# Aspirin compliance for cardiovascular disease and colorectal cancer prevention in the uninsured population

**DOI:** 10.1101/656835

**Authors:** Nina Liu, Adithya Mathews, Justin Swanson, Rahul Mhaskar, Akshay Mathews, Noura Ayoubi, Abu-Sayeef Mirza

**Author notes:** Corresponding author (AM). These authors contributed equally to this work (Nina Liu, Adithya Mathews).

## Abstract

Aspirin is an effective anti-inflammatory and antiplatelet agent as an irreversible inhibitor of cyclooxygenase. In 2016, the U.S. Preventive Services Task Force recommended aspirin for primary prevention of cardiovascular disease (CVD) and colorectal cancer (CRC) in patients aged 50 to 69 with a 10% or greater 10-year CVD risk. Due to the lack of literature describing compliance with these recommendations in the uninsured patient population, we studied the aspirin adherence for CVD and CRC prevention in several free medical clinics. We investigated the records of 8857 uninsured patients who visited nine free medical clinics in the Tampa Bay Area in 2016-2017. Aspirin compliance was assessed for patients with prior myocardial infarction (MI) or coronary artery disease (CAD). 54% (n=1467) of patients met the criteria to take aspirin for primary prevention of CVD and CRC, but just 17% of these patients aged 50-59 were on the medication. 16% percent of patients aged 60-69 were taking aspirin and significantly more men than women were on aspirin (p=0.025). Of the 306 patients who had prior MI or CAD, 50% were on the medication for secondary prevention. Among the uninsured population, there is low compliance with recommendations for aspirin usage to reduce the risk of CVD and CRC. This study demonstrates that further improvements are needed to increase adherence to current guidelines and address barriers uninsured patients may face in maintaining their cardiovascular and colorectal health.

## Introduction

Cardiovascular disease (CVD) is the leading cause of death in the United States according to 2016 data published in the National Vitals Statistics Report from the Centers for Disease Control and Prevention [1]. As an irreversible inhibitor of cyclooxygenase, aspirin is widely prescribed for CVD prevention due to its effectiveness as an anti-inflammatory and antiplatelet agent [2–4]. Long-term use at low doses has additionally been associated with decreased risk of colorectal cancer (CRC) [5]. In 2016, the U.S. Preventative Services Task Force (USPSTF) recommended aspirin for primary prevention of CVD and CRC in patients aged 50 to 69 with a 10% or greater 10-year CVD risk. To determine the 10-year CVD risk, the USPSTF used a calculator derived from the American College of Cardiology and American Heart Association (ACC/AHA) pooled cohort equations. This is a class B recommendation for 50-59 year-olds and a class C recommendation for 60-69 year-olds [6]. Class B recommendations indicate that “there is high certainty that the net benefit is moderate or there is moderate certainty that the net benefit is moderate to substantial”, whereas Class C recommendations indicate “at least moderate certainty that the benefit is small” and to offer the intervention to “selected patients depending on individual circumstances” [7]. Current guidelines from the AHA and American College of Cardiology Foundation (ACCF) recommend aspirin for patients with prior myocardial infarction (MI) or coronary artery disease (CAD) as secondary prevention of CVD [8].

Currently, no studies have examined the use of aspirin in reducing the risk of CVD and CRC in the uninsured population. One study found low rates of aspirin use for primary prevention in low-income, minority populations [9]. Several studies have reported the underutilization of aspirin for primary and secondary CVD prevention among the general population, without distinguishing patients’ insurance status. The suboptimal rates of usage are due to both the lack of physician recommendation to use aspirin and patient non-compliance with physician recommendations [10–12]. Uninsured patients receive less treatment and poorer management of major CVD risk factors than insured patients, despite similar prevalence of disease [13]. The cost of medications is a particularly important factor in managing the health of uninsured patients. The cost-effectiveness of aspirin is well-documented, augmenting its value in managing CVD in uninsured patients [14].

The purpose of this study was to evaluate the compliance with USPSTF and AHA/ACCF recommendations for aspirin use in the primary and secondary prevention of CVD and risk reduction for CRC among uninsured patients. Our secondary purpose was to assess inappropriate aspirin use by uninsured patients who do not meet criteria for its usage.

## Methods

We conducted a retrospective chart review that included 8857 uninsured patients who visited nine free medical clinics in the Tampa Bay Area between January 1, 2016 and December 31, 2017. Age, sex, race, employment status, and history of MI, CAD, cerebrovascular accident, and gastrointestinal ulcers were recorded. Cholesterol levels, blood pressure, diabetic status, and smoking status were used to calculate 10-year Framingham risk scores [15]. The Framingham risk score predicts the risk of developing hard coronary heart disease (myocardial infarction or coronary death) within 10 years. Framingham scores, rather than the ACC/AHA pooled cohort equations, were used to determine 10-year CAD risk because the patient information collected through our chart review did not include all data points required by the pooled cohort equations. The data for total cholesterol was not available, however low density lipoprotein cholesterol levels were available. As a result, Framingham risk scores were more representative of the CVD risk for our population of patients.

Ten-year Framingham risk scores for CAD were calculated for only the 50-69 year-old population to determine which patients qualified to take aspirin for primary prevention of CVD and CRC. Records of patients with a 10% or greater risk score were reviewed to determine if they were taking aspirin for primary prevention of CVD and CRC, and records of patients with prior MI or CAD were reviewed to determine if they were taking aspirin for secondary prevention of CVD.

The secondary aim of this study was to evaluate inappropriate use of aspirin, which we defined as patients taking aspirin with no history of MI, CAD, or cerebrovascular accident, having less than a 10% 10-year Framingham risk score, or having gastrointestinal (GI) ulcers. Although current guidelines from the AHA and American Stroke Association recommend aspirin for patients with a history of ischemic stroke [16], increased risk of bleeding, including history of GI ulcers, is a contraindication for aspirin usage [6]. Given the limited data set, our parameters defining inappropriate aspirin use did not incorporate the various other factors that increase risk of bleeding, such as bleeding disorders, severe liver disease, renal failure, and thrombocytopenia. The data was edited and analyzed using IBM SPSS Statistics software. P-values for categorical data, such as demographics, were determined using Chi-squared and Fisher’s exact test. This study was approved by the University of South Florida Institutional Review Board for the Protection of Human Subjects.

## Results

Of the 8857 patients, 1773 met the USPSTF and AHA/ACCF criteria for taking aspirin. Of the 2724 (30.8%) patients who were 50-69 years old and met the USPSTF criteria, 1467 (53.9%) had a 10% or greater 10-year Framingham risk score, the threshold to take aspirin for primary prevention of CVD and CRC. Three hundred six patients (3.5%) had a history of MI or CAD, meeting AHA/ACCF criteria to use aspirin for secondary prevention of CVD (Fig. 1).

**Fig. 1.**
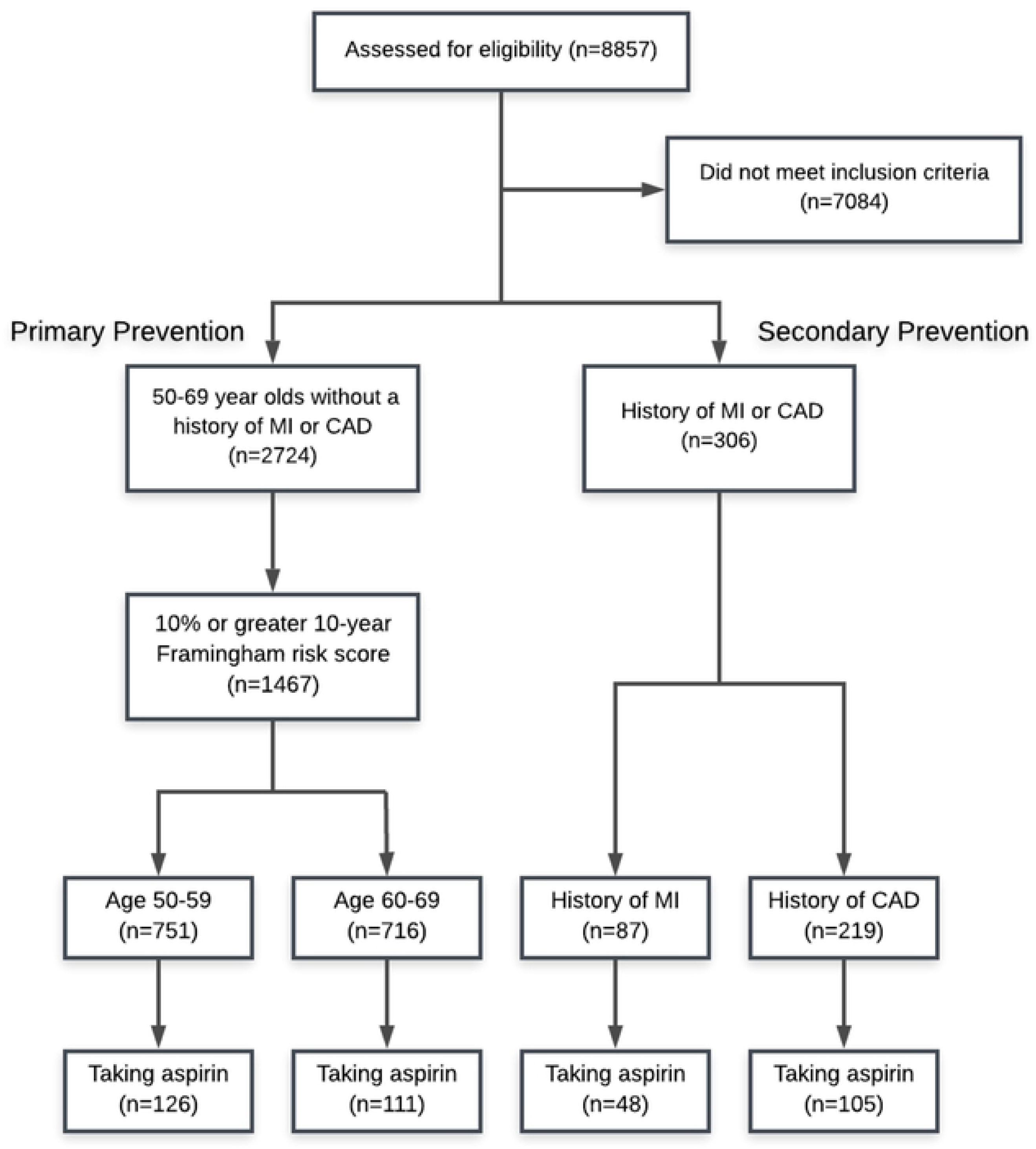
Details of sample development.

Of the 1467 patients who met the criteria to take aspirin for primary prevention of CVD and CRC, 751 were aged 50-59 and 716 were aged 60-69. In the 50-59 year age group, 16.8% (126/751) of the patients were taking aspirin, and sex, race, or employment status did not differ between aspirin users and non-users. In the 60-69 year age group, 15.5% (111/716) were taking aspirin (Fig. 2), significantly more men than women were taking aspirin (p=0.025), and Caucasians were more likely to be taking aspirin than patients of other races (p=0.011) (Table 1).

**Table 1.**
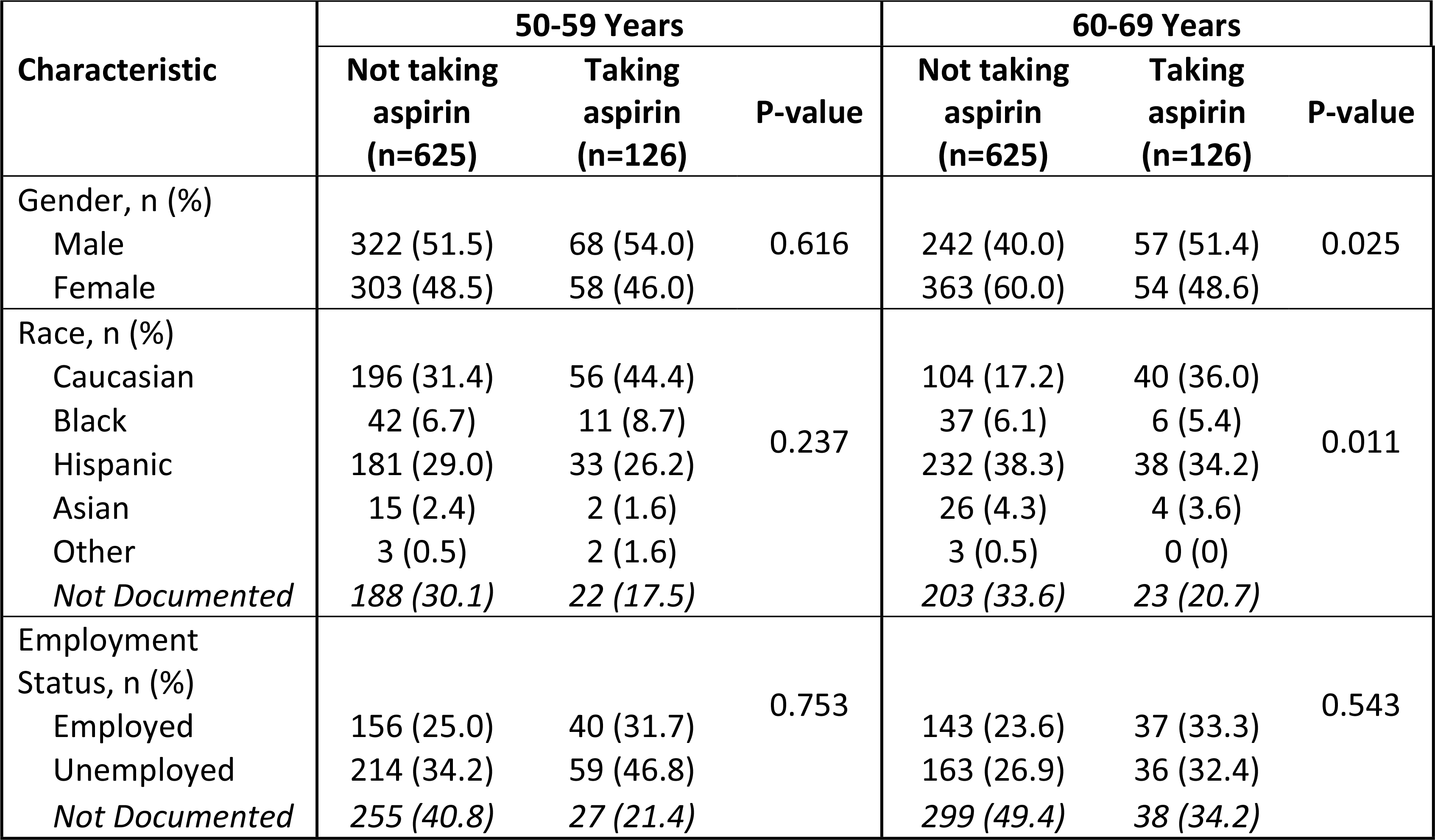
Patient predictors for aspirin use as primary prevention.

| Characteristic | 50-59 Years |  |  | 60-69 Years |  |  |
| --- | --- | --- | --- | --- | --- | --- |
|  | Not taking aspirin (n=625) | Taking aspirin (n=126) | P-value | Not taking aspirin (n=625) | Taking aspirin (n=126) | P-value |
| Gender, n (%) |  |  |  |  |  |  |
| Male | 322 (51.5) | 68 (54.0) | 0.616 | 242 (40.0) | 57 (51.4) | 0.025 |
| Female | 303 (48.5) | 58 (46.0) |  | 363 (60.0) | 54 (48.6) |  |
| Race, n (%) |  |  |  |  |  |  |
| Caucasian | 196 (31.4) | 56 (44.4) | 0.237 | 104 (17.2) | 40 (36.0) | 0.011 |
| Black | 42 (6.7) | 11 (8.7) |  | 37 (6.1) | 6 (5.4) |  |
| Hispanic | 181 (29.0) | 33 (26.2) |  | 232 (38.3) | 38 (34.2) |  |
| Asian | 15 (2.4) | 2 (1.6) |  | 26 (4.3) | 4 (3.6) |  |
| Other | 3 (0.5) | 2 (1.6) |  | 3 (0.5) | 0 (0) |  |
| Not Documented | 188 (30.1) | 22 (17.5) |  | 203 (33.6) | 23 (20.7) |  |
| Employment Status, n (%) |  |  |  |  |  |  |
| Employed | 156 (25.0) | 40 (31.7) | 0.753 | 143 (23.6) | 37 (33.3) | 0.543 |
| Unemployed | 214 (34.2) | 59 (46.8) |  | 163 (26.9) | 36 (32.4) |  |
| Not Documented | 255 (40.8) | 27 (21.4) |  | 299 (49.4) | 38 (34.2) |  |

**Fig. 2.**
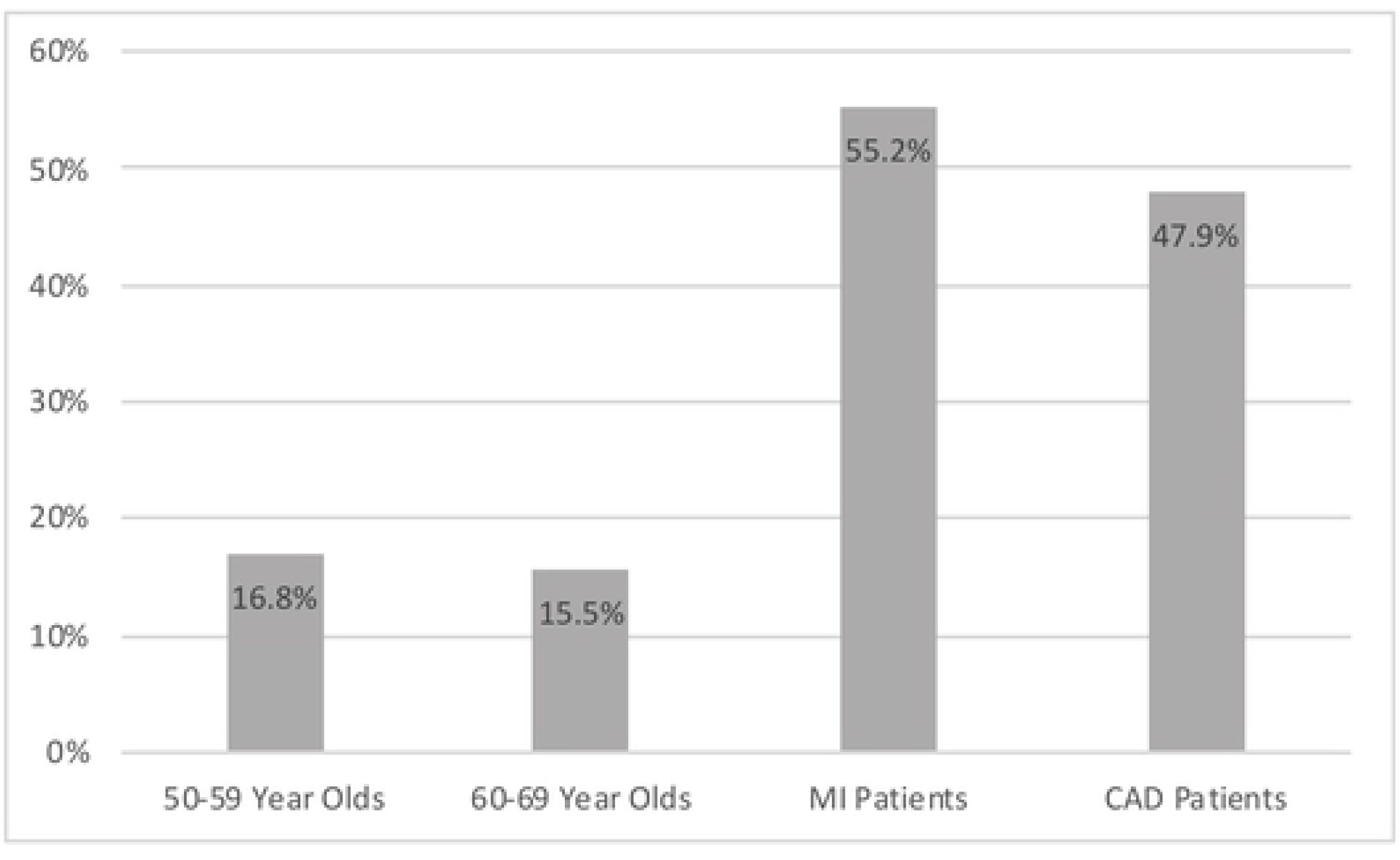
Patients who met criteria and were taking aspirin.

Of the 306 patients who met criteria to take aspirin for secondary prevention of CVD, 50% (153/306) were taking aspirin. 55.2% (48/87) of patients who had a prior MI and 47.9% (105/219) of patients who had a history of CAD were taking aspirin (Fig. 2). The effects of sex and race on compliance with aspirin recommendations were not statistically significant. Among patients with a history of CAD, those who were currently employed were more likely to be taking aspirin (p=0.04) (Table 2).

**Table 2.** Patient predictors for aspirin use as secondary prevention.

| Characteristic | MI Patients |  |  | CAD Patients |  |  |
| --- | --- | --- | --- | --- | --- | --- |
|  | Not taking aspirin (n=39) | Taking aspirin (n=48) | P-value | Not taking aspirin (n=114) | Taking aspirin (n=105) | P-value |
| Gender, n (%) |  |  |  |  |  |  |
| Male | 28 (71.8) | 36 (75.0) | 0.736 | 65 (57.0) | 70 (66.7) | 0.165 |
| Female | 11 (28.2) | 12 (25.0) |  | 48 (42.1) | 35 (33.3) |  |
| Race, n (%) |  |  |  |  |  |  |
| Caucasian | 16 (41.0) | 25 (52.1) | 0.974 | 45 (39.5) | 48 (45.7) | 0.977 |
| Black | 2 (5.1) | 2 (4.2) |  | 11 (9.6) | 11 (10.5) |  |
| Hispanic | 11 (28.2) | 13 (27.1) |  | 30 (26.3) | 26 (24.8) |  |
| Asian | 1 (2.6) | 1 (2.1) |  | 6 (5.3) | 7 (6.7) |  |
| Other | 0 (0) | 0 (0) |  | 0 (0) | 0 (0) |  |
| *Not Documented* | *9 (23.1)* | *7 (14.6)* |  | *22 (19.3)* | *13 (12.4)* |  |
| Employment Status, n (%) |  |  |  |  |  |  |
| Employed | 6 (15.4) | 18 (37.5) | 0.059 | 24 (21.1) | 38 (36.2) | 0.04 |
| Unemployed | 20 (51.3) | 21 (43.8) |  | 48 (42.1) | 38 (36.2) |  |
| *Not Documented* | *13 (33.3)* | *9 (18.8)* |  | *42 (36.8)* | *29 (27.6)* |  |

To assess the inappropriate use of aspirin, we excluded 1467 patients aged 50-69 who had a 10% or greater 10-year Framingham risk score, 306 patients with a history of MI or CAD, and 144 patients with a history of stroke. 172 (2.5%) of the remaining 6919 patients were inappropriately taking aspirin. 6 (6.3%) of 96 patients with history of GI ulcers were taking aspirin, despite an increased risk of bleeding.

## Discussion

Our results demonstrate suboptimal rates of aspirin use in the prevention of CVD and CRC according to current guidelines. Although 53.9% (1467/2724) of uninsured patients met criteria to be on an aspirin regimen for primary prevention, only 16.2% (237/1467) of them were taking the medication. Among uninsured patients with a history of MI or CAD, 50% (153/306) were taking aspirin.

Previous studies reported underutilization of aspirin for the primary and secondary prevention of CVD in the general population. One study reported 40.9% of patients were told by their physician to take aspirin for primary prevention, with 79% complying. Comparably, 75.9% of patients were told by their physician to take aspirin for secondary prevention, with 89.9% complying [12]. Despite the seeming underuse of CVD risk score calculators, such as the Framingham risk score calculator, providers may be considering the risk of GI bleeding and hemorrhagic stroke associated with an aspirin regimen when evaluating patients. A low dose aspirin regimen was found to increase the risk of GI bleeding by 58% and hemorrhagic stroke by 27% in patients using the medication for primary prevention of CVD [17]. Consideration of the bleeding risks could play a role in the under-prescription of aspirin.

Within the general population, aspirin use is lower than that recommended by current guidelines. Our results showed that uninsured patients had even lower rates of use than the general population, indicating room to improve compliance to guidelines. These findings bring up the question of why uninsured patients have suboptimal rates of aspirin use. Lack of health insurance and a low socioeconomic status have been associated with medication non-adherence [18,19]. However, there is limited information regarding provider prescribing patterns in free medical clinics. A combination of poor medication adherence and provider prescribing patterns could be a possible explanation for the discrepancy between aspirin use among the uninsured and general patient populations.

We also found inappropriate use of aspirin. Among patients in our study, 2.5% (172/6919) did not meet guideline criteria but were taking aspirin. This result contrasts with that from a previous study using a national database, which reported that 11.6% of patients were inappropriately taking aspirin for primary prevention of CVD [20]. However, that study used older guideline criteria that defined inappropriate aspirin use as an aspirin regimen in patients with a less than 6% 10-year risk of a CVD event. The 4% difference in guideline criteria could account for the variance in frequency. The association between lack of health insurance and medication non-adherence could also contribute to the lower rate of inappropriate aspirin use among uninsured patients compared to the general population.

The results of our study highlight the need to educate uninsured patients and their providers of the value of aspirin and current clinical guidelines regarding its use. Aspirin is an inexpensive preventive measure, thus it is a valuable and cost-effective tool to prevent CVD and CRC in the uninsured population. The National Heart, Lung, and Blood Institute reported that educational outreach visits, along with audit and feedback strategies, were effective methods to improve clinical practice guideline implementation [21]. These interventions could be used in free medical clinics to address the suboptimal rates of aspirin use. Further studies are needed to improve our understanding of the discrepancy in rates of aspirin use between uninsured patients and the general population. Emphasis should be placed on developing solutions to minimize the gap between these two populations.

The 2019 ACC/AHA Guideline on the Primary Prevention of Cardiovascular Disease recommends considering aspirin for primary prevention in certain patients aged 40-70 who are not at increased risk of bleeding. New studies demonstrating a lack of benefit have influenced the evolving opinion on the beneficial effects of aspirin on primary prevention of CVD [22]. Despite the changing views on aspirin, our study indicates that uninsured patients will still be taking aspirin at lower rates than the general population unless interventions are implemented. This study has several limitations. Patient races were not consistently recorded in medical records, thus we could not fully assess relationships between race and aspirin use. Additionally, our limited dataset did not include comprehensive information regarding factors that can lead to increased risk of bleeding. As a result, it is conceivable that many more patients were inappropriately on low-dose aspirin. The retrospective chart review conducted for this study did not allow us to discern whether physicians discussed potential aspirin regimens with qualified patients. Furthermore, we could not assess if concerns regarding risk of bleeding were addressed in those conversations or if patients decided not to adhere to an aspirin regimen.

In conclusion, patients without health insurance are taking aspirin for the prevention of CVD and CRC at suboptimal rates. Aspirin is underutilized among this population even in comparison to the general population. Further improvements are needed to increase adherence to current guidelines and address barriers uninsured patients may face in maintaining their cardiovascular and colorectal health.

## Acknowledgements

We would like to thank the directors and staff of the nine free medical clinics that allowed us to collect data and make this study possible. We would also like to thank the team of undergraduates, medical students, graduate students, and physicians that contributed to our project.

